# *slendr*: a framework for spatio-temporal population genomic simulations on geographic landscapes

**DOI:** 10.1101/2022.03.20.485041

**Authors:** Martin Petr, Benjamin C. Haller, Peter L. Ralph, Fernando Racimo

**Affiliations:** Lundbeck Foundation GeoGenetics Centre, Globe Institute, University of Copenhagen, Denmark; Section for Molecular Ecology and Evolution, Globe Institute, University of Copenhagen, Denmark; Department of Computational Biology, Cornell University, Ithaca, NY, USA; Institute of Ecology and Evolution, University of Oregon, Eugene, OR, USA

## Abstract

One of the goals of population genetics is to understand how evolutionary forces shape patterns of genetic variation over time. However, because populations evolve across both time and space, most evolutionary processes also have an important spatial component, acting through phenomena such as isolation by distance, local mate choice, or uneven distribution of resources. This spatial dimension is often neglected, partly due to the lack of tools specifically designed for building and evaluating complex spatio-temporal population genetic models. To address this methodological gap, we present a new framework for simulating spatially-explicit genomic data, implemented in a new R package called *slendr* (www.slendr.net), which leverages a SLiM simulation back-end script bundled with the package. With this framework, the users can programmatically and visually encode spatial population ranges and their temporal dynamics (i.e., population displacements, expansions, and contractions) either on real Earth landscapes or on abstract custom maps, and schedule splits and gene-flow events between populations using a straightforward declarative language. Additionally, *slendr* can simulate data from traditional, non-spatial models, either with SLiM or using an alternative built-in coalescent *msprime* back end. Together with its R-idiomatic interface to the *tskit* library for tree-sequence processing and analysis, *slendr* opens up the possibility of performing efficient, reproducible simulations of spatio-temporal genomic data entirely within the R environment, leveraging its wealth of libraries for geospatial data analysis, statistics, and visualization. Here, we present the design of the *slendr* R package and demonstrate its features on several practical example workflows.

## Introduction

Most evolutionary processes in nature have a spatial dimension. Indeed, since its beginnings, the field of population genetics has aspired to build interpretable models of spatial population dynamics (Guillot *et al*., 2009; Barton, Etheridge and Véber, 2013). These include classic theoretical models such as Fisher’s wave-of-advance model (Fisher, 1937), Wright’s isolation-by-distance model (Wright, 1943), Kimura’s stepping-stone model (Kimura, 1953; Kimura and Weiss, 1964), and Malecot’s lattice model (Malécot, 1951; Nagylaki, 1976; Rousset, 1997). The field also has a long history of modeling continuous spatial genetic variation (Levene, 1953; Slatkin, 1973; Barton, 1979; Beerli and Felsenstein, 2001; McRae, 2006; Duforet-Frebourg and Blum, 2014; Bradburd, Coop and Ralph, 2018), inferring spatial covariates associated with genetic patterns (Hanks and Hooten, 2013) and detecting spatial barriers to migration (Safner *et al*., 2011; Petkova, Novembre and Stephens, 2016; Ringbauer *et al*., 2018; Al-Asadi *et al*., 2019; Marcus *et al*., 2021). However, these latter efforts are hampered by a lack of good theoretical predictions for continuous, two-dimensional models (Felsenstein, 1975; Barton, Depaulis and Etheridge, 2002), and simulations can provide a valuable tool in the absence of analytical theory.

The dramatic increase in the number of published whole-genome sequences in the last 20 years (1000 Genomes Project, 2010; Mallick *et al*., 2016; Palkopoulou *et al*., 2018; Feuerborn *et al*., 2021), and the advent of ancient genomics (Green *et al*., 2010; Rasmussen *et al*., 2010), have revealed previously unknown migration events in the history of several species, such as dogs (Bergström *et al*., 2020), horses (Librado *et al*., 2021), elephantids (Meyer *et al*., 2017), and humans (Lazaridis *et al*., 2014; Fu *et al*., 2016). Since migration of populations involves spatial displacement, populations trace their ancestry to different geographic locations (Ralph and Coop, 2013; Osmond and Coop, 2021; Wohns *et al*., 2022). In the context of human history, processes including past migration, gene flow, and population turnovers have been shown to have had a major influence on the present-day distribution of genomic variation (Pickrell and Reich, 2014; Slatkin and Racimo, 2016). Properly anchoring these past demographic events in both time and space has been a focus for new modeling approaches (Racimo *et al*., 2020; Osmond and Coop, 2021; Wohns *et al*., 2022), and is a question of high interest not only in genetics (Bradburd and Ralph, 2019) but also in ecology (Frachetti *et al*., 2017; Loog *et al*., 2017; Crabtree *et al*., 2021; Delser *et al*., 2021).

Despite the key role of geography in population genetics, tools specifically designed for describing and simulating complex spatio-temporal processes are still lacking. Spatial simulations are important not just for rigorous testing and evaluation of existing inference tools and facilitating the development of new inference methods (Liu *et al*., 2006; Currat and Excoffier, 2011; Delser *et al*., 2021; Osmond and Coop, 2021; Wohns *et al*., 2022), but also for gaining intuition about the expected behavior of the processes influencing the patterns of genetic variation under various scenarios of spatial population dynamics (Felsenstein, 1975; Slatkin and Excoffier, 2012). While powerful simulation approaches based on coalescent theory have been developed (Hudson, 2002; Ewing and Hermisson, 2010; Staab *et al*., 2015; Kelleher, Etheridge and McVean, 2016), these have little or no notion of spatiality due to fundamental obstacles to incorporating space into the coalescent framework (Barton, Depaulis and Etheridge, 2002; Barton, Etheridge and Véber, 2010, 2013), although recent algorithmic advances are promising (Kelleher, Etheridge and Barton, 2014). The first pioneering attempt at simulating spatial population genetic data was the software package SPLATCHE (Currat, Ray and Excoffier, 2004; Currat *et al*., 2019). However, SPLATCHE’s simulation engine is limited to discrete demes based on the stepping-stone model, allows simulation of no more than two populations co-existing at a time, and is not suitable for simulating sequence data at a whole-genomic scale (Currat *et al*., 2019). The most advanced simulator with spatial capabilities is currently the forward population genetic simulation framework SLiM (Haller and Messer, 2017, 2019). Highly popular in the population genetics community, SLiM contains a vast library of features for simulating individuals in continuous space (as opposed to older approaches based on discrete demes), including spatial interactions between individuals, neighborhood-based mate selection, and customisable offspring dispersal (Haller and Messer, 2019). Moreover, the recent implementation of tree-sequence recording in SLiM has opened up the possibility of efficient simulation of massive genome-scale and population-scale datasets (Haller *et al*., 2019).

Despite these advances in population genetic simulations, geospatial data analysis remains a complex field with a steep learning curve. Performing even basic manipulations of spatial cartographic objects, handling diverse data formats, and transforming data between different projections and coordinate reference systems (CRS) requires a non-trivial amount of domain-specific knowledge (Lovelace, Nowosad and Muenchow, 2019). Moreover, because the technicalities of geospatial computation are generally not within the scope of population genetic software, available tools do not provide dedicated functionality for building complex and dynamic spatial population models in a straightforward manner. Developing such models and simulating data from them currently requires hundreds of lines of custom code, which is error-prone and hinders reproducibility. Additionally, the lack of specific frameworks for analyzing and visualizing spatially-explicit genomic data further hinders the methodological and empirical progress in spatial population genetics. A flexible and easy-to-use simulation framework specifically designed for developing spatio-temporal population models and analyzing spatial genomic data would expand the horizons of the field, allowing researchers to evaluate the accuracy of novel spatial methods, to test detailed hypotheses about demography and selection, and to answer entirely new kinds of questions about the interactions between organisms across space and time. For instance, many conceptual models and visualizations of past migration events involve depictions of movements of large population ranges across a map as various environmental or cultural conditions change; however, there is currently no easy way to simulate these movements and generate realistic spatio-temporal genomic data.

To address these issues, we have developed a new programming framework, called *slendr*, designed for simulating and analyzing spatially-explicit genomic data (available at www.slendr.net with extensive documentation and tutorials). The core component of this framework is an R package which leverages real Earth cartographic data (or, alternatively, an abstract user-defined spatial landscape) to programmatically and visually encode spatial population boundaries and their temporal dynamics across time and space, including expansions, migrations, population splits, and gene flow. Because of the challenges involved in testing and validating complex models, *slendr* encourages an interactive workflow in which each component of the model can be inspected and visualized as the model is incrementally constructed in a “bottom-up” fashion. Spatio-temporal models programmed in *slendr* can then be executed using a SLiM back-end script which is bundled with the package and can be controlled by a dedicated R function without leaving the R environment. Additionally, traditional, random-mating, discrete-deme, non-spatial population models can also be simulated, either in forward time using the aforementioned SLiM script or using an alternative coalescent *msprime* (Baumdicker *et al*., 2022) back-end script which is also bundled with the R package and can provide a more efficient simulation engine for non-spatial models. Both simulation engines of *slendr* save genomic outputs in the form of an efficient tree-sequence data structure (Kelleher *et al*., 2018), and the *slendr* R package provides a set of functions for loading and processing tree-sequence output files and computing population statistics on them by seamlessly integrating the *tskit* tree-sequence analysis Python module into its R interface. Additional functionality includes conversion of individual trees to a standard R *ape* phylogenetic format (Paradis and Schliep, 2019), and automatic transformation of spatial tree-sequence table data to the standardized *sf* format for geospatial data analysis in R (Pebesma, 2018).

Overall, the *slendr* R package facilitates reproducibility by providing a unified framework for writing complete spatial simulation and analysis pipelines entirely in R, which we demonstrate with several concrete examples.

## Overview of the *slendr* design and typical workflow

From a software design perspective, the *slendr* R package represents a tight integration of three distinct parts. First, it implements an interactive and visually-focused R interface for encoding spatio-temporal population dynamics focused on building arbitrarily complex models from small individual components (i.e., simple R objects), designed to require only a minimum amount of code. Second, *slendr* includes two back-end simulation scripts implemented in SLiM (Haller and Messer, 2019) and *msprime* (Baumdicker *et al*., 2022). These scripts are bundled with *slendr*, are specifically tailored to interpret *slendr* demographic models, and produce tree-sequence files as output (Haller *et al*., 2019). Lastly, *slendr* provides an interface to the *tskit* tree-sequence analysis library (Kelleher *et al*., 2018). Although this library is written in C and Python, *slendr* exposes its functionality to the R environment in an R-idiomatic way, blending it naturally with the popular *“tidyverse”* philosophy of data analysis (Wickham *et al*., 2019).

Although these three parts operate at fundamentally different levels under the hood, this integrated approach allows all steps of a *slendr* workflow—from specifying spatio-temporal demographic models, to executing simulations and analyzing simulation results—to be performed without leaving the R environment (**Figure 1**). This allows the user to leverage R’s features for visualization and interactive data analysis at every step of the analytic pipeline, and facilitates reproducibility by eliminating the need to manually integrate disparate software tools and programming languages (Sandve *et al*., 2013). In this way, *slendr* follows the footsteps of the original design of the S (and later R) languages: to present a consistent and convenient data-analysis–focused domain-specific front end to more efficient and faster tools written in other languages and frameworks (in this case SLiM and *msprime*) (Chambers, 2020).

**Figure 1.**
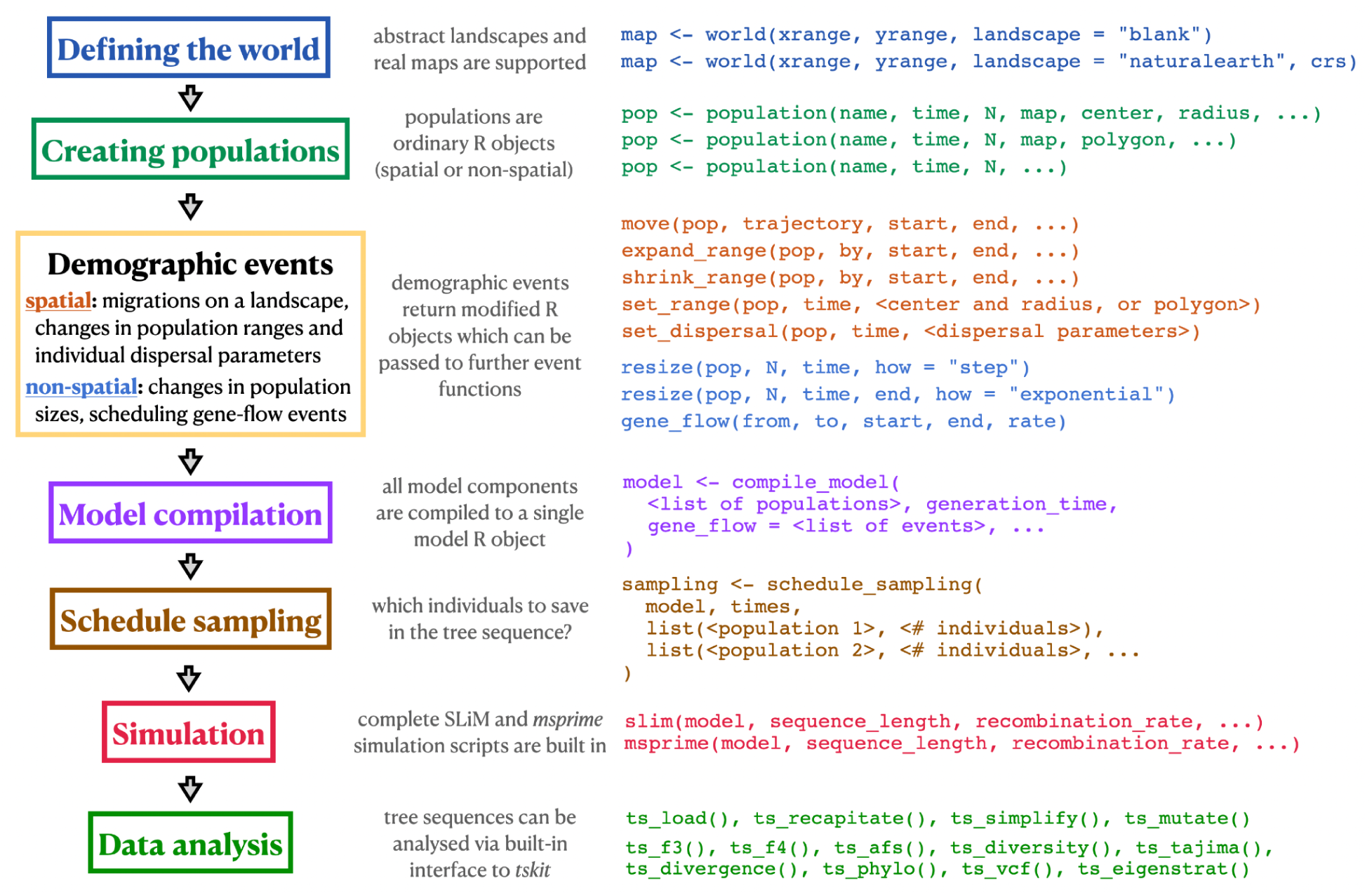
Schematic overview of a hypothetical *slendr* simulation and analysis workflow. The colored rectangles on the left indicate individual steps of a hypothetical *slendr* workflow. Short code snippets in matching colors on the right show examples of *slendr*’s declarative interface used in each step, focusing only on a selected few relevant functions and their most important arguments (additional optional arguments are replaced by the “…” ellipsis symbol). The full function reference index can be found at slendr.net/reference. Note that regardless of whether a spatial or non-spatial *slendr* model is being defined and simulated, the workflow remains identical: the same functions are used for both types of models, and the spatial or non-spatial nature of a model is automatically detected by *slendr*.

In the remainder of this section we outline the individual steps of a typical *slendr* simulation and analysis workflow, as well as describe the individual building blocks of the three main components of the *slendr* framework mentioned above.

## Defining the world

At the beginning of a *slendr* workflow, the user defines the parameters of the world that the simulation will occupy using the function `world()` (**Figure 1**). If the simulated world represents a region on Earth, the appropriate set of vectorised spatial features will be automatically downloaded from a public-domain cartographic database (www.naturalearthdata.com). The user can also specify a dedicated coordinate reference system (CRS) appropriate for the projection of the geographic region of interest in order to minimize the distortion of distances and shapes inherent to transforming geometries (in this case population ranges and landscape features) from the three-dimensional Earth surface to its two-dimensional representation on a map. Alternatively, the world can be represented by an abstract landscape, optionally with custom features such as islands, barriers, or corridors. If a non-spatial deme-based model is to be simulated, this step can be omitted and no changes to the downstream steps described below are needed.

## Creating populations and scheduling demographic events

Populations in *slendr* are created with the `population()` function which creates a simple R object containing the parameters of the population that was created (**Figure 1**). In addition to specifying the name, time of appearance, and initial number of individuals for the new population, the user can also specify a `world` object and, if desired, a set of coordinates for the spatial range that the population will occupy. For convenience, the coordinates of all spatial objects in *slendr* (maps, geographic regions, population ranges) are always specified in the global geographical CRS (i.e., degrees of longitude and latitude) but are then automatically internally transformed into the chosen projected CRS (which uses units of meters) if it was specified when creating the `world` (**Figure 1**). This way, users can encode spatial coordinates in familiar units of longitude and latitude while *slendr* internally maintains the proper shapes and distances of spatial features by performing all spatial transformations in the projected CRS.

All *slendr* spatial objects are internally represented using a data type implemented by the R package *sf* (Pebesma, 2018), which has emerged as the *de facto* standard for geospatial data analysis in R (Lovelace, Nowosad and Muenchow, 2019). Despite the convenience of the *sf* framework, manipulation of geospatial objects in *sf* still requires writing a non-trivial amount of code dealing with low-level technical details (manipulating and transforming the coordinates of points, lines and polygons). Because most of these technical details are not relevant for specifying population genetic models, we designed a set of domain-specific functions for encoding spatial population dynamics which are expressed in terms of population genetics concepts rather than geometric transformations (**Figure 1**). For instance, the `move()` function accepts a *slendr* population object (i.e., internally an `sf` object, encapsulating the low-level geometric coordinates of the population), a trajectory given as a list of coordinates in longitude and latitude, and a timespan over which the population displacement should occur (*Example 3* and **Figure 4**). Other kinds of dynamic spatial events (population range expansions and contractions, for example) are implemented in an analogous manner. Other demographic events, such as population size changes and gene flow, can be scheduled similarly with another set of straightforward functions (**Figure 1**).

For spatial models, the user has the option to fine-tune the within-population individual dispersal and mating dynamics (described in detail in *Example 2* and **Figure 3**) using a set of parameters such as the maximum mating distance between individuals, the dispersal distance of offspring from their parents (and the kernel function of this dispersal), or the parameter influencing the uniformity of the dispersal of individuals within their population’s spatial boundary. These can be assigned for each population separately or kept at their default values given in the `compile_model()` step (as we show in *Example 2*). The `competition` parameter determines the maximum neighborhood distance in which individuals in a SLiM simulation compete with each other for space. If this distance is small, then individuals with nearby neighbors have much lower fitness. If the distance is larger, then the effects of crowding are more diffuse. However, if this distance is larger than the dispersal distance (as in *Example 2*), populations tend to self-organize into an evenly-spaced grid of patches. (**Figure 3C**). Using the `competition` parameter, within-population dynamics can thus be fine-tuned to represent various levels of individual clustering into sub-groups (**Figure 3C**). In addition to the competition parameter, a `mating` parameter determines the maximum distance to which an individual will look for a mate to produce offspring. Finally, a `dispersal` parameter determines how far an offspring can end up from its parent, and a related `dispersal_fun` argument characterizes the density function for this dispersal: “normal” (default), “uniform”, “cauchy”, “exponential”, or “brownian”; more details are available in the *slendr* R package documentation at slendr.net/reference. We note that changes in all three spatial interaction and dispersal parameters can be also scheduled dynamically at specific times throughout the run of a model with a *slendr* function `set_dispersal()`.

A standard feature of many population genetic frameworks is the specification of the times of various demographic events in terms of generations, either forwards in time starting from generation 1 (as is the case with SLiM) or backwards in time starting from time 0 “in the present’’ (as is the case with coalescent frameworks such as *msprime*). This can be cumbersome in cases when the events or samples of interest are traditionally specified in times of “years before present” (such as dated ancient DNA samples), or in situations in which it would be desirable to simulate future outcomes, as in ecological predictive modeling. Moreover, because these standard times often need to be converted into generations by a factor specifying the length of the generation time of the species of interest, this can easily lead to frustrating bugs in simulation scripts. To ameliorate this situation, *slendr* allows the users to specify times in whichever time units they would prefer, in either the forward or backward direction. The time direction is automatically detected by *slendr* from the sequence of demographic events specified for a model (but can also be set explicitly), and the conversion of event times into generations is performed in the compilation step via the provided `generation_time` argument to `compile_model()` (described below). Similarly, times of the tree-sequence nodes in *slendr*’s outputs (which are specified by most simulation software in terms of generations backwards in time) are automatically converted by *slendr* back into the units of time used by the user during model specification.

Because every *slendr* demographic event function returns a modified population object which can be further used as an input to other *slendr* functions, the R interface encourages a workflow in which complex models are composed incrementally from smaller components (**Figure 1**, *Examples 1-3*). Importantly, because each *slendr* function assures the consistency of the model by enforcing appropriate constraints during the model definition process (e.g., a population cannot be moved or participate in a gene-flow event at a time when it would not yet exist), this workflow facilitates the early discovery of bugs before the simulation (which can be extremely computationally costly) is even executed. This is further facilitated by a convenient set of plotting functions, such as `plot_map()` and `plot_model()`, which can visualize the spatio-temporal dynamics of the specified model (or its individual components) as the model is being incrementally developed.

## Model compilation

Having defined all the individual components of a population model (i.e., created all the necessary population and gene-flow events), the user calls the function `compile_model()` to compile the model configuration to a single R object (**Figure 1**)—a step in which *slendr* performs additional checks for model consistency and correctness. Furthermore, this operation also transforms the model components from their R representation into a set of files on disk, written in a format interpretable by the built-in SLiM and *msprime* simulation back-end scripts which are used to execute *slendr* models in the next phase, as described below. The compiled model object can also be used as input for a built-in R-based interactive browser app built using the *shiny* R package (Chang *et al*., 2021) which allows the user to “play” the defined spatial model dynamics over time and explore the “admixture graph” implied by the model (Patterson *et al*., 2012) for additional verification of the model’s correctness. The functions `plot_map()` and `plot_model()` mentioned above also accept a compiled model object as their input and produce a static visualization of the model.

## Scheduling sampling events and simulation

The *slendr* package comes bundled with two simulation back-end scripts which were tailored to interpret the configuration files produced by the compile_model() function and simulate the model, triggering all of the encoded population dynamics in the course of the simulation run.

The first back-end script is written in SLiM’s programming language Eidos (Haller and Messer, 2019), and can execute both spatial and non-spatial *slendr* models in a Wright–Fisher setting by calling *slendr*’s `slim()` function. The second back-end script is implemented using *msprime* (Baumdicker *et al*., 2022) and is designed to interpret the compiled *slendr* model in a non-spatial setting as a standard coalescent simulation by calling *slendr*’s `msprime()` function. Both simulation engines can interpret the same *slendr* model without a need to make any changes. For instance, a spatial model can be run with the *msprime* back end, in which case the spatial component of the model is simply ignored. Because coalescent simulations are generally much more computationally efficient than their forward-time counterparts, the *msprime* back end of *slendr* can be useful for R users who would like to run a large number of traditional, non-spatial simulation replicates efficiently without having to write custom Python *msprime* code or use its *ms*-like command-line interface (Hudson, 2002). Importantly, the correctness of both *slendr* simulation engines is validated using a set of automatic statistical tests on non-spatial models which ensure that when a *slendr* model is run in both SLiM and *msprime*, the demographic events specified by the model (population splits, population size changes, and gene-flow events) result in equivalent site-frequency spectra and *f*-statistics (Patterson *et al*., 2012) between both back ends.

Leveraging the ability to save simulation outputs as a tree sequence (Kelleher *et al*., 2019; Speidel *et al*., 2019) from both SLiM (Haller *et al*., 2019) and *msprime* (Baumdicker *et al*., 2022), *slendr* embraces the tree sequence as its primary output format. This is powerful not only because the tree sequence represents an extremely efficient representation of even large-scale population genomic data, but also because it provides an elegant way to calculate many population genetic statistics of interest, a feature which we describe in more detail in the next section. To specify which simulated individuals should be recorded in the output tree sequence, *slendr* provides two alternative approaches. First, if no explicit sampling schedule is specified, all individuals living at the very end of a SLiM simulation run are explicitly sampled (i.e., “remembered”) in the tree sequence output, matching the default behavior of SLiM. If a *slendr* model is simulated with the *msprime* back end, the number of recorded individuals will be equal to the population size of each population at the start of the coalescent process looking backwards in time (i.e., in “the present”). Alternatively, *slendr* provides a flexible way to trigger sampling events via its `schedule_sampling()` function, which allows one to specify the time (and, optionally, the location) at which a sample comprising a given number of individuals from a given population should be taken and recorded in the tree sequence (*Example 3*). To improve readability and interpretation of *slendr* analysis code, every sampled individual can be referred to using its readable name during tree-sequence processing and computation of statistics (*Examples 1, 2, and 4)* rather than just by numeric identifiers as is the case with the default tree-sequence analysis workflow with *tskit* (Kelleher *et al*., 2018).

## Data analysis

The default output of a *slendr* simulation is a tree sequence. However, because processing and analysis of tree-sequence files requires a non-trivial knowledge of Python or C (Kelleher *et al*., 2018) which many R users might not have, *slendr* provides an R-idiomatic interface to the most commonly used *tskit* tree-sequence methods such as the allele frequency spectrum, Patterson’s *f*-statistics, and various summary statistics of population diversity (Patterson *et al*., 2012; Ralph, Thornton and Kelleher, 2020). This way, users can design population genetic models in R, execute them from R using the built-in `slim()` or `msprime()` functions, and analyze the resulting tree sequence data without having to leave the R environment for downstream statistical analyses and plotting, and without the need to convert outputs to other bioinformatic or population genetic file formats. Although primarily designed for analysis of tree sequences generated from *slendr* models, the R-*tskit* interface can operate also on tree sequences without *slendr*-specific metadata. Therefore, users who would prefer to run simulations with standard *msprime* or SLiM scripts but are interested in analyzing their tree-sequence results in R will still find the *slendr* R package useful. The reference manual at slendr.net/reference contains a complete list of *tskit* tree-sequence methods that have been integrated into *slendr*’s R interface. If integration with traditional tools such as PLINK (Purcell *et al*., 2007) or ADMIXTOOLS (Patterson *et al*., 2012) is required, functions for exporting to VCF (Danecek *et al*., 2011) and EIGENSTRAT (Patterson *et al*., 2012) are also provided.

During a spatial simulation in SLiM, each sampled individual’s location on the simulated landscape is tracked and recorded in the tree sequence, encapsulating the full spatio-temporal genealogical history that has been simulated. When the tree-sequence output file is then loaded by *slendr*, *slendr* processes the spatial locations of nodes in the tree sequence (which represent chromosomes of past and present individuals), and transforms them back into the original coordinate system of the simulated world, adding additional annotation data such as readable names of sampled individuals, population assignments of each individual and node, etc. Furthermore, this information is exposed in an *sf*-compatible format, meaning that the spatio-temporal information about ancestral relationships between simulated samples can be processed, analyzed, and visualized using a wide range of R packages including *sf*, *ggplot2,* and *dplyr* (Pebesma, 2018; Wickham *et al*., 2019). Additionally, individual trees in the tree sequence can be extracted by a *slendr* function, `ts_phylo()`, which converts *tskit*-formatted tree objects into the format defined by the R phylogenetics package *ape*, which has been the standard for phylogenetics in the R ecosystem for nearly two decades (Paradis and Schliep, 2019). This gives *slendr* users even more options to analyze tree-sequence results with a large array of standard phylogenetics tools available for the R environment (Paradis, 2011).

## Installation and software dependencies

*slendr* is currently developed for macOS and Linux. It is available on the CRAN R package repository at https://CRAN.R-project.org/package=slendr, and can be installed from the interactive R console with the standard command `install.packages(”slendr”)`. Development versions of *slendr* which contain latest bug fixes and new experimental features can be installed from its GitHub repository using the R package *devtools* with the R command `devtools::install_github(“bodkan/slendr”)`.

Two external software dependencies must be present on a user’s system to leverage the full functionality of *slendr*: a forward population genetic simulator SLiM (Haller and Messer, 2019) (which is required for running spatial simulations and non-spatial simulations in the forward-time setting) and a trio of Python modules *msprime* (Baumdicker *et al*., 2022), *tskit* (Kelleher *et al*., 2018) and *pyslim* (github.com/tskit-dev/pyslim) (which are needed to run *slendr* models as coalescent simulations and to analyze tree-sequence data).

The SLiM software is available for all major operating systems and its installation instructions can be found at messerlab.org/slim. Importantly, the current version of *slendr* requires the latest release of SLiM 4.0. In order to use SLiM for simulations in *slendr*, the R session needs to be aware of the path to the directory containing the SLiM binary. Calling `library(slendr)` for the first time provides an informative message for the user on how this can be accomplished by modifying the `$PATH` variable by editing the `∼/.Renviron` file.

Because some users might find the experience of setting up a dedicated Python environment with the necessary Python modules challenging (especially users who exclusively work with R), *slendr* provides an R function `setup_env()` which automatically downloads a completely separate Python distribution and installs the required versions of *tskit*, *msprime*, and *pyslim* Python modules in their correct required versions into a dedicated virtual environment without any need for user intervention. Moreover, this Python installation and virtual environment are isolated from other Python configurations that might be already present on the user’s system, thus avoiding potential conflicts with the versions of Python and Python modules required by *slendr*. Once this isolated Python environment is created by `setup_env()`, users can activate it in future R sessions by calling a helper function `init_env()` after loading *slendr* via `library(slendr)`. Therefore, although *slendr* uses Python modules for internal handling of tree-sequence data and coalescent simulation, direct interaction with Python is not necessary.

## Relationship of *slendr* to SLiM and *msprime*

Given that *slendr*’s simulation engines are implemented in SLiM and *msprime*, it is worth elaborating on its relationship to these simulation frameworks, particularly in terms of the features supported by *slendr*. First, it is important to note that *slendr* is not simply a wrapper for SLiM and *msprime* in the strict sense of the word, since *slendr* does not provide an R equivalent of every function and method provided by SLiM and *msprime*. Instead, *slendr* aims to provide a user-friendly, R-idiomatic way to encode a particular class of “traditional” Wright-Fisher population genetic models frequently used in evolutionary biology and population genetics, allowing users to employ such models with a minimal amount of coding. Most importantly, *slendr* models currently assume that populations evolve via random mating, and that the genomes of individuals evolve neutrally, with mutations overlaid on top of the simulated genealogies after each simulation run. This applies also to spatial *slendr* demographic models, with the caveat that interaction and dispersal distance parameters can—depending on the exact parametrization of each spatial *slendr* model—cause individuals to only mate locally, which can have interesting implications for the behavior of standard population genetic statistics (as shown in *Example 2*).

The complete set of models supported by *slendr* is likely to slightly expand over time as new features are implemented. Details of new features, such as customized recombination maps and non-neutral mutation types, are being discussed with the community on the GitHub page of *slendr* (https://github.com/bodkan/slendr), and users are encouraged to provide feedback there. The four practical examples (*Examples 1–4* below) have been designed to demonstrate the full range of *slendr*’s features at the time of writing.

Finally, because *slendr*’s forward and coalescent simulation back ends are implemented as fairly standard SLiM and *msprime* scripts, the performance of *slendr* simulations and tree-sequence analyses can be assessed using already-existing benchmarks and guidelines provided by publications describing SLiM and *msprime* (Haller *et al*., 2019; Baumdicker *et al*., 2022; Haller and Messer, 2022).

## Practical examples

In the following sections, we present the features of the *slendr* R package with several practical examples, each of which focuses on a different aspect of the *slendr* simulation framework. We start by showing how traditional, non-spatial, random-mating models can be specified with a minimum amount of R code (*Example 1*). We then proceed with two examples of spatial models: first, a model showing how the degree of the spatial spread of a population can be adjusted by setting the within-population individual-based dispersal dynamics (*Example 2*); second, a model which schedules the movements of entire population ranges across a landscape (*Example 3*). These examples are intended to demonstrate *slendr’*s ability to define complex spatio-temporal models incrementally, building them from simpler components. We also emphasize how *slendr* model configuration and simulation steps naturally flow into data analysis, all within the R environment. In the final demonstration (*Example 4*), we tap into the rich information embedded in spatial tree sequences to visualize individual trees on a landscape, tracing the complex spatio-temporal ancestry of an individual on the simulated map. Extended versions of these and many other examples with complete reproducible code for simulation, analysis, and plotting can be found as standard R package vignettes at *slendr*’s website (www.slendr.net).

### Example 1: Traditional non-spatial model

Regardless of whether a spatial or non-spatial model is defined and simulated, the *slendr* workflow remains the same. Therefore, before we explore spatial models, we begin by showing how a traditional, non-spatial population genetic model can be constructed with *slendr* and how users can compute population genetic statistics on simulated tree-sequence outputs using *slendr*’s R interface to the *tskit* tree-sequence analysis library (Kelleher *et al*., 2018) (represented by functions with the `ts_*()` prefix, **Figure 1**).

First, we define an abstract demographic model similar to that which is commonly used in teaching the principles behind the *f*_4_-ratio ancestry proportion estimator (Patterson *et al*., 2012). In *slendr*, we define the model with a straightforward sequence of `population()` calls that schedule the order of splits for several populations, taking care of parent–daughter population relationships by providing the appropriate population object as a `parent` argument when creating each daughter population (**Figure 2A**). We then schedule a single gene-flow event between the populations *“b”* and *“x1”* by calling the `gene_flow()` function. After compiling the model with `compile_model()`, we verify its correctness by visualizing the embedded population relationships with `plot_model()` (**Figure 2B**). Although only a single `gene_flow()` event is featured in this example, more complex gene-flow networks can be specified with *slendr*. Conveniently, strict consistency checks validate each encoded gene-flow event before the computationally costly simulation is run. Examples of complex models with dozens or hundreds of gene-flow events can be found in the documentation available on the *slendr* website (www.slendr.net).

**Figure 2.**
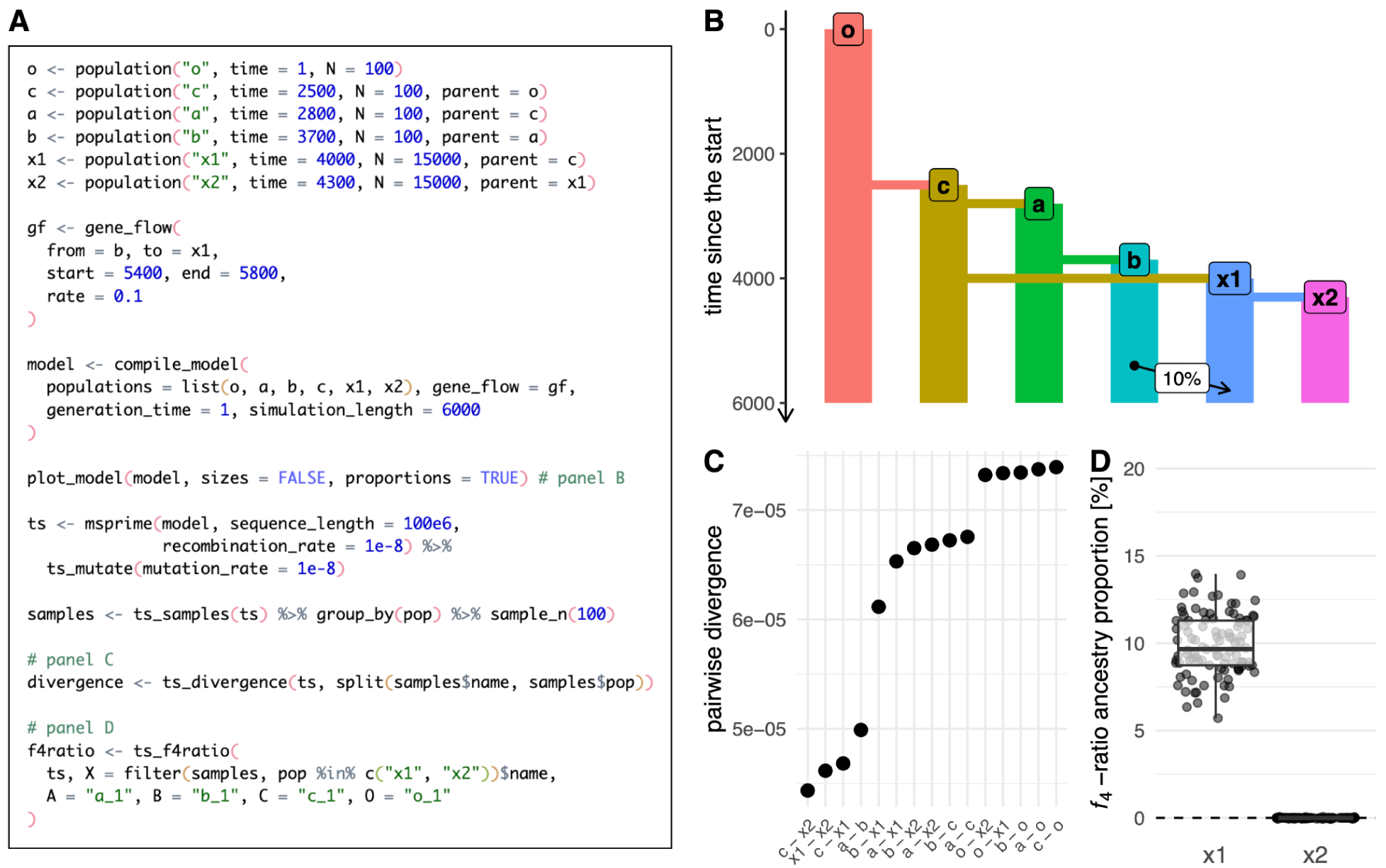
Example 1: specifying a non-spatial model and computing statistics on tree-sequence output. (A) A script which defines a model of a simple demographic history of six populations, simulates it with the *msprime* back end by calling the function `msprime()`, and performs analyses shown in **B**–**D**. **(B)** A visual overview of the compiled *slendr* model produced by `plot_model()` prior to simulation. **(C)** Visualization of the data frame produced by `ts_divergence()` on the output tree sequence simulated. **(D)** Ancestry proportions estimated with `ts_f4ratio()` directly from the tree sequence output. As expected from the model definition, the *f*_4_-ratio statistic estimates indicate ∼10% ancestry from “b” in the population “x1”, but 0% ancestry in population “x2”; this agrees with the model overview shown in panel B. Full *ggplot2* visualization code for the figures can be found in a vignette dedicated to this paper at www.slendr.net. The runtime for the simulation and analysis shown in **A** was ∼5 minutes, as measured on a 16’’ MacBook Pro (2021) equipped with the Apple M1 Pro chip, 32 GB RAM, and running macOS Ventura 13.1.

As stated before, *slendr* provides two simulation back ends; here we use the coalescent *msprime* back end to simulate the model, since SLiM’s spatial capabilities are not required for this simple non-spatial model. However, we note that the function `slim()` could be used in place of the `msprime()` call to perform the equivalent forward-time simulation just as easily. By default, *slendr* automatically loads the simulated tree-sequence object which can be immediately used for analysis. In this example, we compute the pairwise divergence between random samples of 100 individuals from each population with the function `ts_divergence()` (**Figure 2C**). Finally, we use the function `ts_f4ratio()` to compute the values of the *f*_4_-ratioestimate of *“b”* ancestry in populations *“x1”* and *“x2”*, which differ in whether or not they experienced gene flow from *“b”* (**Figure 2D**). All other tree sequence analysis functions of *slendr* (**Figure 1**) can be accessed in the same way. We note that because *slendr* assigns symbolic, permanent names to individuals during sampling, the users can refer to them with these names during tree-sequence operations such as simplification and when computing tree-sequence statistics.

### Example 2: Model with population dispersal dynamics

In our second example, we move from a non-spatial, random-mating model to a model which is explicitly spatial. First, we create an abstract, circular world map using the function `world()`, producing a completely featureless landscape (see *Example 3* for a more elaborate world map). We then create a series of eight populations which all occupy that map, as specified by the `map` argument to `population()`, but do not interact with each other. For simplicity, each population forms its own evolutionary lineage without additional splits or gene-flow events. Importantly, we set the `competition` parameter of each population to a value which forces the individuals to assume an increasing degree of spatial subdivision which, in turn, affects the amount of diversity expected in each population. Finally, we compile the model to a single object with `compile_model()` and run it with the `slim()` back end, simulating 16.000 diploid genomes of 10 megabases each (**Figure 3A**). After the simulation finishes, we simplify the produced tree sequence, overlay mutations on the simulated genealogies, and use the *slendr* function `ts_diversity()` to compute the expected heterozygosity in a sample of 100 individuals from each population, inspecting how heterozygosity is affected by the emergent spatial arrangement of each population (**Figure 3B, C**). We note that some of the values of the spatial competition distance parameter used in this example are quite large, especially compared to the much shorter maximum distance of individual dispersal and mating. Although biologically rather unrealistic, the competition distances have been chosen to give rise to very different degrees of spatial subdivision and, consequently, to varying levels of population genetic diversity, with the intention to demonstrate the ease with which a wide range of model dynamics can be configured by the user.

**Figure 3.**
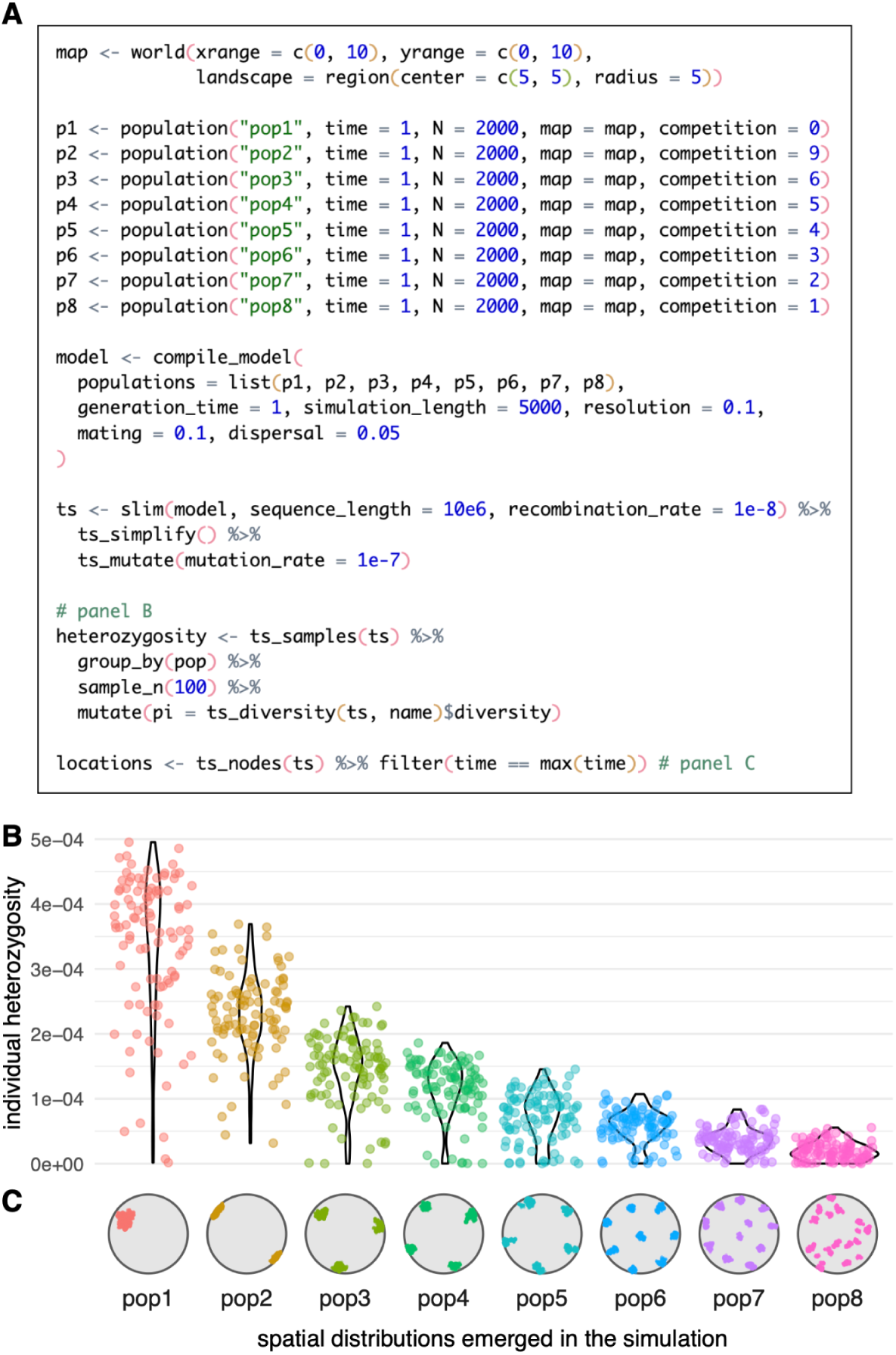
Example 2: a spatial model which involves the parametrization of within-population dispersal dynamics. (A) A complete script which defines eight populations as independent lineages or species, each with constant size and each defined with a different value of *slendr*’s spatial competition parameter, with analysis code to produce panels **B**–**C**. The simulation is run with *slendr*’s SLiM back end for 5000 generations, after which a tree sequence recording the genealogical history of 2000×8 diploid individuals is loaded, simplified, and mutated. Heterozygosity is then computed for 100 individuals randomly sampled from each population at the end of simulation. **(B)** Distribution of heterozygosities of individuals observed in all eight populations. **(C)** A snapshot of the spatial distributions which emerged as a result of the `competition` parameter value set for each population. Full visualization code for the figures can be found in a vignette dedicated to this paper at www.slendr.net. The runtime for the simulation and analysis shown in **A** was ∼12 minutes, as measured on a 16’’ MacBook Pro (2021) equipped with the Apple M1 Pro chip, 32 GB RAM, and running macOS Ventura 13.1.

### Example 3: A toy model of movements and expansions of human populations in West Eurasia over the last 50,000 years

In this example we further expand on the *slendr* functionality demonstrated in the first two examples, introducing programming of expansions and migrations of entire population ranges across a realistic landscape—perhaps the most distinctive feature of *slendr*. The model we implemented here is inspired by large-scale population migrations and turnover events inferred from ancient DNA analyses of human remains from across West Eurasia (Lazaridis *et al*., 2014; Allentoft *et al*., 2015; Haak *et al*., 2015), although we caution that it is an extremely simplified toy model intended only as an illustrative example.

Similarly to *Example 2*, we begin by defining a world map for the simulation (**Figure 4A**), in this case using realistic Earth cartographic data provided by the Natural Earth project (naturalearthdata.com). Because we focus on the broad region of West Eurasia, we select the most appropriate coordinate reference system (CRS) for projecting this region on a two dimensional map which is EPSG:3035. We then define a series of populations, specifying their approximate geographic ranges using simple polygons. We then use the functions `move()` and `expand_range()` to schedule when and where populations should migrate, and by what distance and how quickly their population ranges should expand across the landscape during simulation. We again use `plot_model()` to visualize the demographic history embedded in the *slendr* model as a non-spatial tree-like structure with gene-flow edges (**Figure 4B**); here, we also use `plot_map()` to get a “compressed” overview of the spatio-temporal population range dynamics on the simulated map (**Figure 4C**). We note that unlike in the two previous examples, which were specified in forward time units, this example expresses the timing of demographic events in units of “years before present” which is more natural to this model.

**Figure 4.**
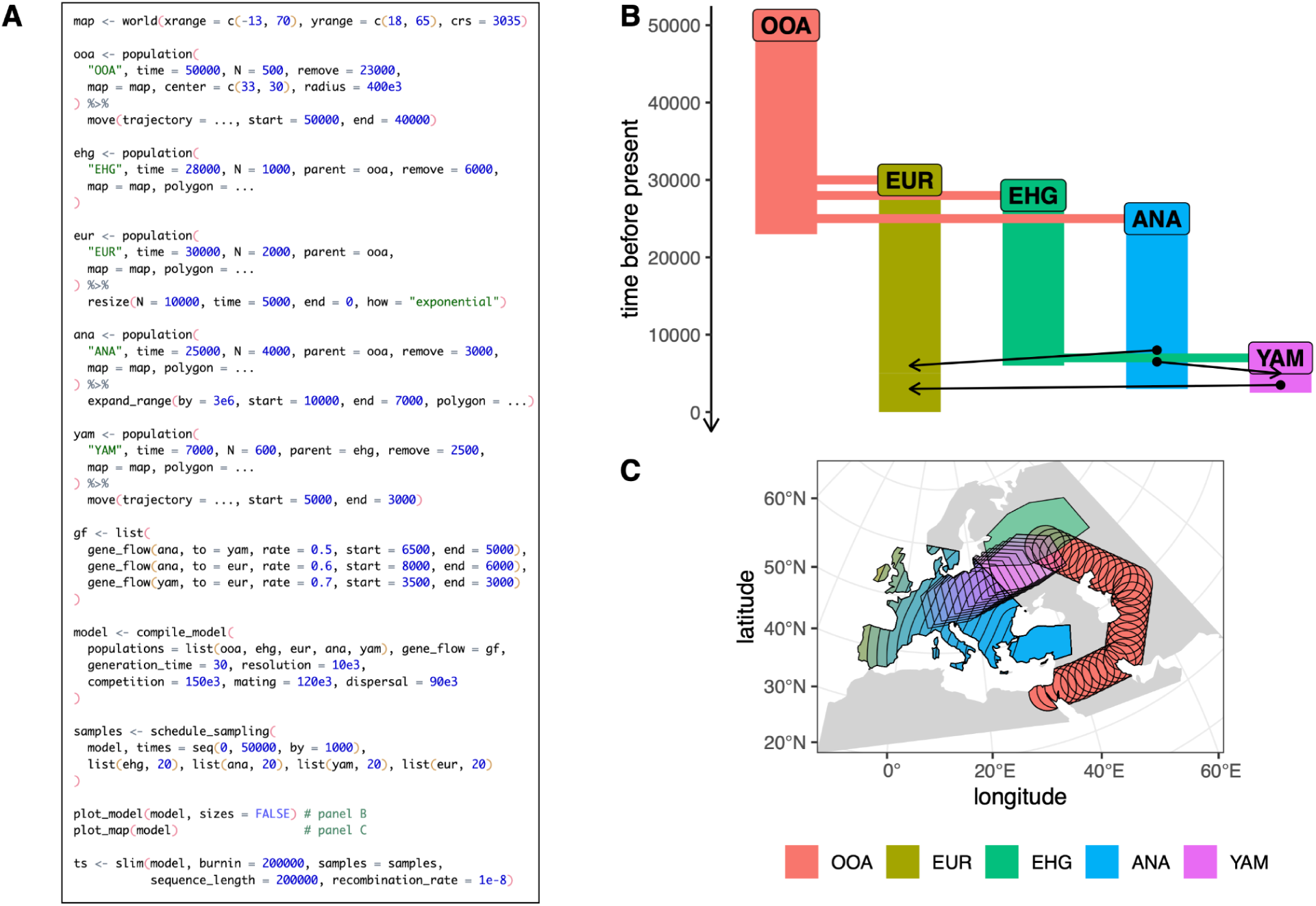
Example 3: a demographic model on a real Earth landscape. (A) A *slendr* script which defines a toy spatio-temporal model of human prehistory in West Eurasia, with analysis code that produces panels **B**–**C**. For brevity, we do not specify the full set of coordinates for each spatial demographic event or population range polygon, instead indicating them as `”…”`; the complete reproducible code can be found in a vignette dedicated to this paper at www.slendr.net. **(B)** Visual summary of the non-spatial component of the demographic model, produced by `plot_model()` with arrows indicating gene flow events. **(C)** A “compressed” view of spatio-temporal snapshots of population ranges throughout the course of the model prior to the simulation, produced by `plot_map()`. The runtime for the simulation shown in **A** was ∼3 minutes, as measured on a 16’’ MacBook Pro (2021) equipped with the Apple M1 Pro chip, 32 GB RAM, and running macOS Ventura 13.1.

In the previous two code examples (**Figure 2A, 3A**) we used the default tree-sequence sampling of *slendr*, which implicitly records the genomes of all the diploid individuals alive at the end of a simulation. In this example, we instead use `schedule_sampling()` to specify a series of sampling events from each population every 1,000 years. We then execute the compiled model and the sampling schedule specified using the `slim()` back-end, which records only the scheduled set of sampled individuals in the tree-sequence output file.

### Example 4: Visualization of individual trees and spatio-temporal ancestral lineages across a landscape

In our final example (**Figure 5**), we return to the abstract toy model of West Eurasian prehistory developed in *Example 3*. To leverage *slendr*’s power to simulate genomic data from complex spatial demographic models, *slendr* makes it easy to tap into the large library of geospatial data science packages available for R (Lovelace, Nowosad and Muenchow, 2019) by automatically converting simulated spatial locations to an *sf*-compatible tabular format (Pebesma, 2018), as we will see here.

**Figure 5.**
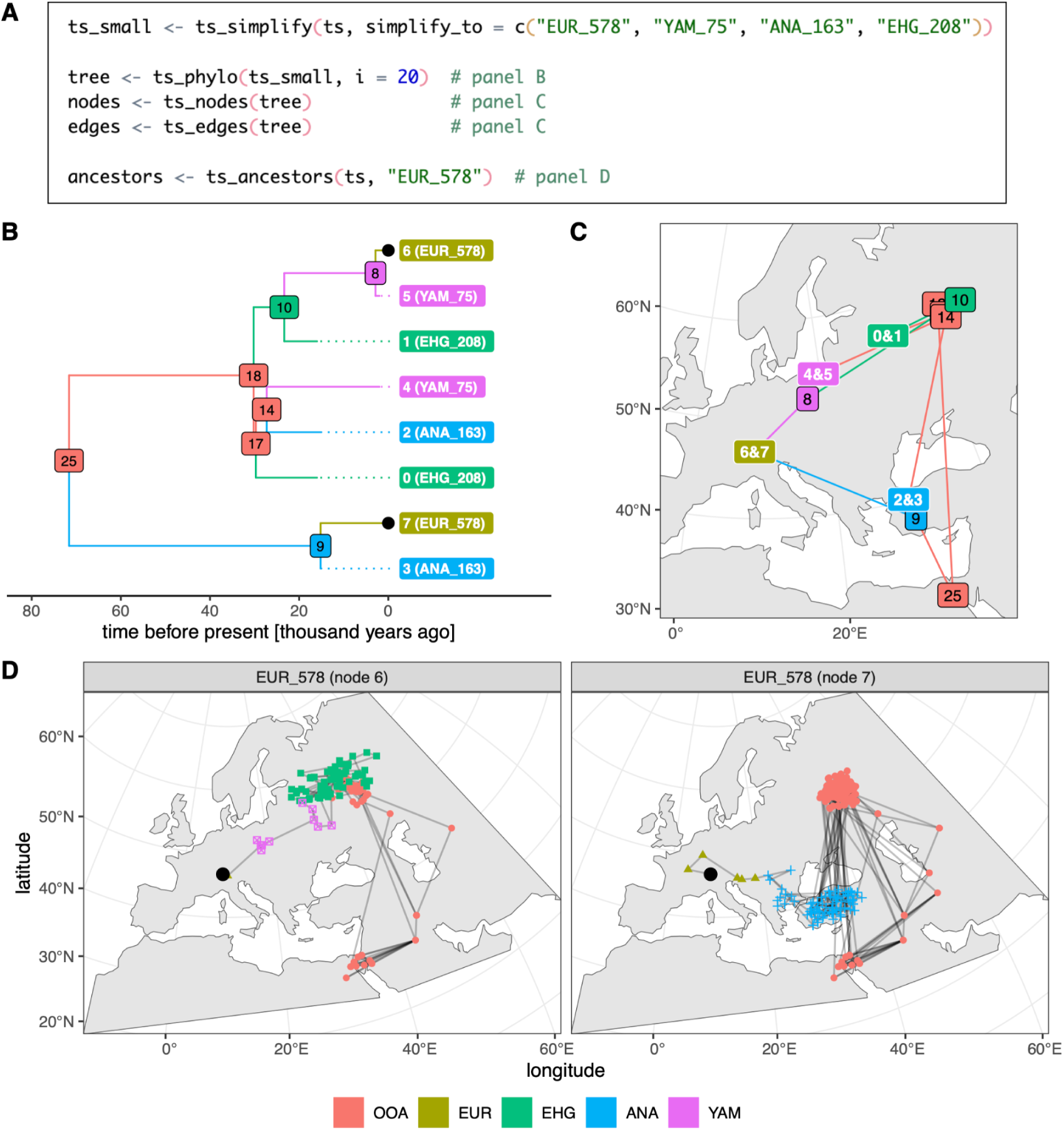
Example 4: accessing and visualizing spatio-temporal information encoded in trees and tree sequences simulated with the *slendr*. (A) A continuation of the script from Example 3, showing how a (potentially very large) tree sequence generated from a *slendr* model can be simplified to a subset of individuals with `ts_simplify()`. A single tree from the tree sequence is then extracted with `ts_phylo()`, the tables of spatio-temporal locations of nodes and branches of the tree are extracted by `ts_nodes()` and `ts_edges()`, and ancestry information for one individual across the entire tree sequence is extracted with `ts_ancestors()`, in order to produce the data plotted in **B**–**D**. **(B)** A visualization of the tree extracted by `ts_phylo()` using standard visualization features of the *ggplot2* and *ggtree* R packages. Dotted lines indicate shortened branches of ancient samples. **(C)** Visualization of the tree from panel **B** as a network across the original spatial simulation landscape, with each node indicating the location of a particular individual who lived at some point during the simulation. Labels with two numbers correspond to the locations of sampled individuals, each carrying two chromosomes which are represented by two nodes in the tree sequence. All node numbers correspond to those shown in the tree in panel **B**. The plot was generated with *ggplot2* using the *sf*-formatted data extracted by `ts_nodes()` and `ts_edges()`. **(D)** A visualization of the spatio-temporal ancestry of a single simulated European individual, “EUR_578”, using the information from the entire tree sequence. Each sub-panel shows the spatial ancestry distribution of one of the two chromosomes carried by this individual (the location of whom is indicated by a black dot), tracing its ancestry through different lineages all the way back to a population in Africa. For easier reference, the same black dots indicate the two chromosomes of this individual also in the tree in panel **B**. The *ggplot2* code for the figures is omitted for brevity. Full reproducible code examples including the visualization code can be found in a vignette dedicated to this paper at www.slendr.net. The runtime for the code shown in **A** was ∼1 second, as measured on a 16’’ MacBook Pro (2021) equipped with the Apple M1 Pro chip and 32 GB RAM, running macOS Ventura Version 13.1.

To demonstrate the richness of the spatio-temporal information recorded in the tree sequence, we use the full tree sequence produced by the code in **Figure 4A** and simplify it so that it contains only the history of a small subset of the thousands of individuals sampled during the spatio-temporal simulation (**Figure 5A**). We then extract the 20th tree in the tree sequence with *slendr*’s function `ts_phylo()`, which converts a tree from the *tskit* tree sequence into an R `phylo` format defined by the *ape* R package, a standard tool for phylogenetics in R (Paradis and Schliep, 2019). Such tree objects can be analyzed by any of the dozens of R packages which operate on *ape*’s `phylo` trees—for instance, in **Figure 5B** we show a visualization of this tree using the R package *ggtree* (Yu *et al*., 2017). Furthermore, because the tree was generated from a spatially-annotated tree sequence, the user can extract information about the location of each individual (or node) in the tree across space and time, as well as ancestral relationships between nodes in the tree, using `ts_nodes()` and `ts_edges()` respectively. Crucially, because these functions automatically convert locations into *sf*’s geospatial representation (including the appropriate CRS projection), the results can be immediately plotted on a map with *ggplot2*, which has built-in support for *sf* data (**Figure 5C**).

In addition to extracting and visualizing single trees representing a genealogy of a set of sampled genomes descending from a common ancestor (spatial or non-spatial), *slendr* also provides a way to extract the complete spatio-temporal ancestry of a single sample going back in time across the entire tree sequence, potentially spanning many trees with thousands of the sample’s ancestors. This can be accomplished with the function `ts_ancestors()` which, in an analogous way to `ts_nodes()` and `ts_edges()`, exposes the spatio-temporal information in the tree sequence as an *sf* object which can be visualized on a map with *ggplot2*. In this example, we use `ts_ancestors()` to reconstruct the spatio-temporal ancestry distribution for a single simulated European individual (“EUR_578”, represented by the black dot in **Figure 5D**). Because this individual is diploid, we can trace the ancestry carried by its one chromosome through an expansion from Anatolia (**Figure 5D**, right panel), while its other chromosome clearly traces its ancestry to a population which migrated to central Europe from an eastern population (**Figure 5D**, left panel).

Note that by default, the tree sequence output of a *slendr* simulation only contains information about ancestors which are represented by coalescent nodes in some marginal tree—i.e., nodes which are a most recent common ancestor of some pair of sampled nodes. In this example, in **Figure 5B** and **C** we can see that the most immediate ancestor (node number 9) of one chromosome of the sampled individual “EUR_578” lived in the region of Anatolia, but the ancestor of its second chromosome lived in Europe (node number 8); but we do not know where all the ancestors along the edges between nodes 9-7 and 8-6, since they were simplified away. Similarly, **Figure 5D** shows the distributions of locations of most recent common ancestors, not all ancestors. The distribution of ancestors at a particular point in time could be obtained by adding an appropriate sampling event to *slendr*’s sampling schedule and then extracting ancestors from that time.

## Discussion

The *slendr* R package provides a new programmable framework for simulating complex spatio-temporal genomic data. The package implements a set of features for defining spatial population range dynamics with a declarative and visually-focused R interface and uses a tailor-made SLiM script as an efficient population genetic simulation engine. Additionally, *slendr* provides a convenient new way to simulate and analyze large-scale genomic data sets even from traditional, non-spatial demographic models using *msprime* entirely within the R environment.

Owing to its declarative interface, which requires little code even for complex models, the *slendr* package is highly accessible even to researchers or students with little or no prior experience in programming. One of the major challenges for novice population geneticists is having to learn how to integrate multiple different software tools and programming frameworks. R (R Core Team, 2021) is often the first language that biology and bioinformatics students learn, since it offers a large number of libraries for data analysis, statistics, and plotting (Wickham and Grolemund, 2016). For these users, *slendr* provides the opportunity to explore population genetic concepts and simulate realistic population genomic data as soon as they learn the most basic principles of R (i.e., how to call R functions and work with data frames), without first having to learn Python for *msprime* simulations (Baumdicker *et al*., 2022), shell scripting for simulators from the *ms* family (Hudson, 2002; Staab *et al*., 2015), or Eidos for SLiM (Haller and Messer, 2019).

Tree sequences provide an efficient way to compute many commonly used population genetic statistics directly on the simulated genealogies (Ralph, Thornton and Kelleher, 2020); because *slendr* uses the tree sequence as its default output format (Kelleher *et al*., 2018; Haller *et al*., 2019), in many cases users do not need to convert simulation outputs to external file formats such as VCF or EIGENSTRAT for analysis in other software. This way, *slendr* simulations can be readily used in model fitting and population genetic analyses in situations which have traditionally required converting simulated data to genotype files before analyzing them with population genetics tools such as PLINK (Purcell *et al*., 2007) or ADMIXTOOLS (Patterson *et al*., 2012). That said, export to VCF and EIGENSTRAT genotype file formats is supported with a single function call (`ts_vcf()` and `ts_eigenstrat()`) if needed.

A key principle in the design of *slendr* has been reproducibility (Sandve *et al*., 2013): a complete *slendr* simulation and analysis workflow can be written as a single R script. Additionally, the compilation of any *slendr* module produces a self-contained “bundle directory” containing all model configuration files and simulation back-end scripts required to execute the model from the command-line. Although accessing this directory is not necessary for standard workflows because *slendr* operates entirely from R, these bundles can be checked into a git history and provided as supplementary files along with a publication, allowing independent replication even without relying on *slendr* itself.

Moving forward, we expect that the *slendr* framework will become a useful tool to produce ground-truth data for comparing and benchmarking inference methods for modeling spatial genomic processes (Peter and Slatkin, 2013; Petkova, Novembre and Stephens, 2016; Marcus *et al*., 2021; Muktupavela *et al*., 2022), as well as for the development of new approaches to spatial problems in population genomics. There is great potential for deploying *slendr* in simulation-based inference methods, like Approximate Bayesian Computation (ABC) (Beaumont, Zhang and Balding, 2002; Csilléry *et al*., 2010), thanks to its tight integration with the rest of the R modeling landscape. A major challenge in ABC is the significant amount of coding needed to program simulations of demographic history and integrate them with software for computing population genetic statistics. *slendr* can program complex models and compute relevant statistics using its tree-sequence interface with a relatively small amount of code, all within a single R workflow. Furthermore, although *slendr* does not currently include features for implicit, automated parallelism (an important aspect of computation-heavy modeling approaches such as ABC), users can rely on numerous R packages providing a wide range of parallelization techniques (Eddelbuettel, 2021).

Nonetheless, inference of spatial dynamics from genetic data remains an open research problem with many potential pitfalls, and we strongly caution users to avoid overinterpretation. For instance, *slendr* models retain a notion of discretely delineated populations, but even a reasonable fit of such a model to real data does not erase the reality that such groupings are rarely, if ever, as stable and cleanly distinguished as in idealized models. Indeed, confounding the simple models used in population genetics with reality can be actively harmful (Coop, 2022; Khan *et al*., 2022). Furthermore, population genetic modeling in general is notoriously challenging due to the many parameters involved (Gravel *et al*., 2011; Pickrell and Pritchard, 2012; Kamm *et al*., 2020). In this respect, advanced, explicitly spatial models of the kind unlocked by *slendr* present an even bigger challenge. For instance, how can we best do model comparison, and among what set of models? What would constitute a good “null hypothesis” when modeling potentially complex spatial population dynamics? Furthermore, even relatively simple models can be ill-posed or even nonidentifiable: many combinations of spatial parameters (such as individual dispersal or mating distances) may give rise to similar genetic patterns. As every demographic inference study makes assumptions about the process which generated the data, sometimes explicitly and sometimes implicitly, awareness of these assumptions and careful approach to modeling (Beaumont, 2010; Gerbault *et al*., 2014; Loog, 2021) are vital for correct interpretation of results. We hope that the ease with which *slendr* allows one to explore the impact of spatio-temporal parameters on population dynamics—and the fact that *slendr* forces the researcher to state those parameters explicitly—will help guide researchers in establishing guidelines for good practice, to delineate the limits of what can be learned and, consequently, avoid overinterpretation (or misinterpretation) of such parameters.

In its current version (v0.5.0 as available on the CRAN repository), *slendr*’s spatial simulation maps are limited to landscapes that exhibit binary habitability—i.e., any given location either is or is not habitable by individuals. A more ecologically realistic simulation could allow for varying degrees of habitability at different locations, which would affect the size of the simulated population. Future extensions of the *slendr* framework could include the incorporation of fine-scaled geographic maps storing individual habitability values for each pixel of the raster, allowing for dynamic changes of such maps over time. This would effectively make the size of the population an emergent consequence of the habitability metric aggregated across the map. This extension would require significant changes to the *slendr* back-end code, moving to modeling population densities per unit of landscape area using non-Wright–Fisher dynamics, but the necessary software building blocks are already supported by SLiM and examples of these types of simulations are discussed in the SLiM manual (Haller and Messer, 2022). A recently published Python module *Geonomics* provides an interface for simulating genetic data on arbitrary landscape rasters (Terasaki Hart, Bishop and Wang, 2021). Implementing such functionality in *slendr* would have the advantage of using a much more efficient SLiM simulation engine and a greater ease of use due to *slendr*’s emphasis on visually-focused interactive model design in R. The main challenge would therefore lie in making sure that the additional complexity involved in making the *slendr*’s SLiM back end more flexible does not compromise the current simplicity of its declarative interface. The benefits of this extension would be numerous, including for genomic forecasting and predicting species ranges in the face of climate change and ecological breakdown (Fitzpatrick and Keller, 2015; Exposito-Alonso *et al*., 2019; Theodoridis *et al*., 2020), and for constructing models of species distribution dynamics in the ancient past (Wang *et al*., 2021). Implementation of this extension of *slendr* is still in the planning stages, in collaboration with the community on the project’s GitHub page.

At the moment, slendr can only produce genome sequences from a single species (although with an arbitrary number and spatial arrangement of populations of the species) due to the restrictions imposed by its simulation back end. However, many types of genomic resources distributed across space and time are represented by fragmentary mixtures of genomes from multiple species. This includes ancient microbiomes from human remains (Rasmussen *et al*., 2015), sedimentary DNA from permafrost, caves, or lake and marine cores (Willerslev *et al*., 2003; Parducci *et al*., 2017; Armbrecht *et al*., 2019; Vernot *et al*., 2021), and environmental DNA from water, soil, or air samples (Taberlet *et al*., 2012; Stat *et al*., 2017; Lynggaard *et al*., 2022). Recent developments in SLiM would allow *slendr* to perform multi-species simulations, which would facilitate ecological modeling of species distributions (Fordham *et al*., 2021) or of past epidemics (Duchene *et al*., 2020) from a fully genomic perspective.

Finally, at the time of writing, *slendr* models are limited to neutral simulations, and this restriction applies even to simulations performed via its SLiM back end. In particular, *slendr* does not currently provide built-in support for specifying mutation types, genomic element types, recombination maps, or custom SLiM callbacks. Providing an R equivalent for SLiM’s complete functionality would be a daunting task of limited utility, and would substantially complicate *slendr*’s intuitive R syntax for encoding demographic models (Figure 1). An attractive alternative for supporting more advanced, customized models could be to retain the behavior of *slendr* described in this manuscript as the default, but provide the possibility of overriding different aspects of this behavior by injecting user-defined SLiM snippets at appropriate locations in *slendr*’s SLiM back-end code. We are exploring this possibility for future versions of the software.

Ultimately, we hope that our new simulation framework will help generate new ideas about the insights that can be gleaned from the rich spatio-temporal information hidden within DNA sequences. Furthermore, we aspire to help budding researchers in population genetics get started with simulations and build their intuition about population genetic concepts by developing models using more traditional non-spatial methods and statistics, and we believe that *slendr* could be a useful tool for teaching population genetics to students. We hope that by easily generating and visualizing genomic models on real landscapes, we can spark new ways of thinking about how organisms evolve (Bradburd and Ralph, 2019) and enable clearer discussions about the fundamental interconnectedness of genomes across space and time (Mathieson and Scally, 2020).

## Acknowledgements

We thank Isabel M. Pötzsch, Mariadaria Kathrine Ianni-Ravn, Emma Prantoni, and Moisès Coll Macià for testing and feedback on early versions of slendr, and the members of the Racimo group for valuable comments on the design of the software and this manuscript. FR was supported by a Villum Young Investigator Grant (project no. 00025300), a Novo Nordisk Fonden Data Science Ascending Investigator Award (NNF22OC0076816) and by the European Research Council (ERC) under the European Union’s Horizon Europe programme (grant agreements No. 101077592 and 951385). MP was supported by the aforementioned Novo Nordisk Fonden award, as well as a Lundbeck Foundation grant (R302-2018-2155) and a Novo Nordisk Foundation grant (NNF18SA0035006) given to the Lundbeck GeoGenetics Centre. PR was supported by NIH award R01HG010774.

